# Tracking attention using RIFT with a consumer-monitor setup

**DOI:** 10.1101/2025.10.03.680199

**Authors:** Kabir Arora, Ü. Gülce Çelik, Lasse Dietz, J. Leon Kenemans, Stefan Van der Stigchel, Surya Gayet, Samson Chota

## Abstract

Rapid Invisible Frequency Tagging (RIFT) is a recent technique that extends the traditional frequency tagging approach by modulating the luminance of distinct locations on a screen at frequencies beyond the threshold of perception (≥60Hz). Retrieving the corresponding frequencies from M/EEG recordings provides a time-resolved and location-specific measure of early visual processing, without the confounding effect of introducing visible stimuli. This method is most frequently harnessed as a tracker of covert attention in experimental paradigms across various disciplines in cognitive neuroscience. However, almost all existing RIFT work so far has made use of expensive display hardware limited in its accessibility. Recent work has shown that a RIFT response elicited by a 480Hz refresh-rate consumer monitor can be retrieved from the EEG, but it is not yet clear whether such a setup can be utilized to track the locus of attention. Here, using a spatial cueing paradigm (n=24) while simultaneously tagging two locations (60.0Hz and 65.5Hz), we show that covert attention can be reliably measured with RIFT on a 360Hz refresh rate consumer monitor. We hope that this study will facilitate the widespread application of using consumer-grade high refresh-rate gaming monitors with RIFT for future scientific and applied research.

## Introduction

Frequency tagging is a common stimulation technique where different stimuli are flickered at unique frequencies, and the neural response to these can later be disentangled and uniquely tracked over time. Recent advancements in display technology have led to the emergence of a novel approach to frequency tagging known as Rapid Invisible Frequency Tagging (RIFT) (Zhigalov et al., 2019). Unlike traditional Steady-State Visual Evoked Potential (SSVEP) approaches that use slower flicker frequencies (<30Hz) which can be consciously perceived, RIFT employs a form of rapid stimulation using luminance flickers at frequencies beyond the threshold of perceptibility (≥ 60Hz). RIFT therefore offers a measure of early visual processing at specific locations in the visual field without the involvement of downstream visual processing regions associated with conscious perception (Arora, Gayet, et al., 2025; Drijvers et al., 2021; Seijdel et al., 2023; Zhigalov et al., 2019). RIFT is most frequently utilized as a continuous and “invisible” tracker of spatial attention, measuring how attention influences cortical responses to specific locations in the visual field without (or before) presenting a perceptible stimulus (Arora, Gayet, et al., 2025; Bouwkamp et al., 2025; Zhigalov et al., 2019). However, technical factors limit the feasibility of implementing RIFT in various lab settings. Though RIFT was initially paired with MEG, it has more recently been shown to work with EEG as well (Arora et al., 2024; Dietz et al., 2025; Dimigen et al., 2025; Hustá et al., 2025; Wang et al., 2026), representing a first step in expanding its accessibility. One remaining obstacle, however, is that of the associated display technology.

To date, most RIFT studies have utilized the PROPixx DLP-LED projector for the projection of stimuli on the screen to achieve the high refresh rate required for high-contrast luminance modulation. However, such specialized high-quality projection systems are often prohibitively expensive, which limits their accessibility to research groups. Moreover, the complexity of the setup of such display equipment can further restrict its availability to research laboratories. Conveniently, some recent consumer-grade (gaming) monitors are also able to reach high refresh rates similar to those that have been used with the PROPixx. Since these monitors are available at only a small fraction of the PROPixx’s price, they offer a substantial reduction in hardware costs and provide a cost-effective alternative for conducting RIFT research.

Dimigen et al. (2025) recently utilized a commercially available 480 Hz organic light-emitting diode (OLED) monitor in combination with EEG and showed that such consumer-grade monitors can reliably deliver 480 Hz RIFT stimulation with accurate frame timings, with only rare single missed frames that do not seem to impact visibility (Dimigen et al., 2025). This study demonstrates the presence of a reliable RIFT response at two location conditions (i.e., central and peripheral). However, since this study did not incorporate an attentional manipulation, the essential question of whether RIFT responses measured with such monitors are sensitive to attentional modulations during cognitive experiments remains unanswered. Furthermore, they found that the tagging response from a peripheral stimulus was considerably weaker than that of a central stimulus, leaving open the question of whether tagging responses from peripheral stimuli –as are common in experiments on visuospatial attention –are robust enough to measure modulations of the RIFT response by attention even on a monitor setup.

This project builds on Dimigen et al. (2025) by including a spatial cueing task (to manipulate the allocation of attention) and tagging a larger set of locations in more participants. We had two main aims. Firstly, we utilized an off-the-shelf consumer-grade monitor (i.e., Alienware 500Hz Gaming Monitor AW2524HF) at a refresh rate of 360Hz and aimed to show that a robust RIFT response can be elicited using this novel display setup. Secondly, we examined whether the setup is sensitive enough to capture attentional modulations of the RIFT response across the visual field. To this end, we employed a cueing paradigm in 24 participants to investigate the deployment of covert spatial attention using RIFT in combination with EEG. Specifically, we tested whether a robust RIFT signal could be evoked using a consumer-grade gaming monitor at 360Hz, and crucially whether this would allow us to measure covert attentional modulations across the visual field. We believe that this would mark a key step both in making RIFT research more accessible, and also in allowing for the integration of an extra attentional tracker into pre-existing cognitive tasks at little additional cost.

## Methods

### Participants

We recruited 24 healthy participants (17 female, mean = 22.50 ± 1.69 years) with normal or corrected-to-normal vision. None of the participants reported a history of epilepsy or psychiatric diagnosis. Participants received compensation of either €20 or a corresponding amount of participation credits in line with Utrecht University’s internal participation framework (SONA). The study was conducted in accordance with the protocol approved by the Ethics Committee of the Faculty of Social and Behavioral Sciences at Utrecht University with reference number 863732 –HOMEOSTASIS (approved on 18/03/2020).

### Experimental Design and Procedure

Participants reported the orientation of a target presented at one of eight locations as indicated by a spatial cue (Figure 1). Throughout the trial, a fixation cross was presented centrally, and participants were instructed to keep their gaze fixated on it. The task presented eight identical circles arranged around a central fixation cross in two concentric rings: an inner ring of four circles and an outer ring of four circles positioned further away. The circles served as placeholder target/distractor locations. On each trial, the circle in which the target was presented and the circle which was diagonally opposite to this were invisibly tagged (See *Tagging Manipulation*).

**Figure 1:**
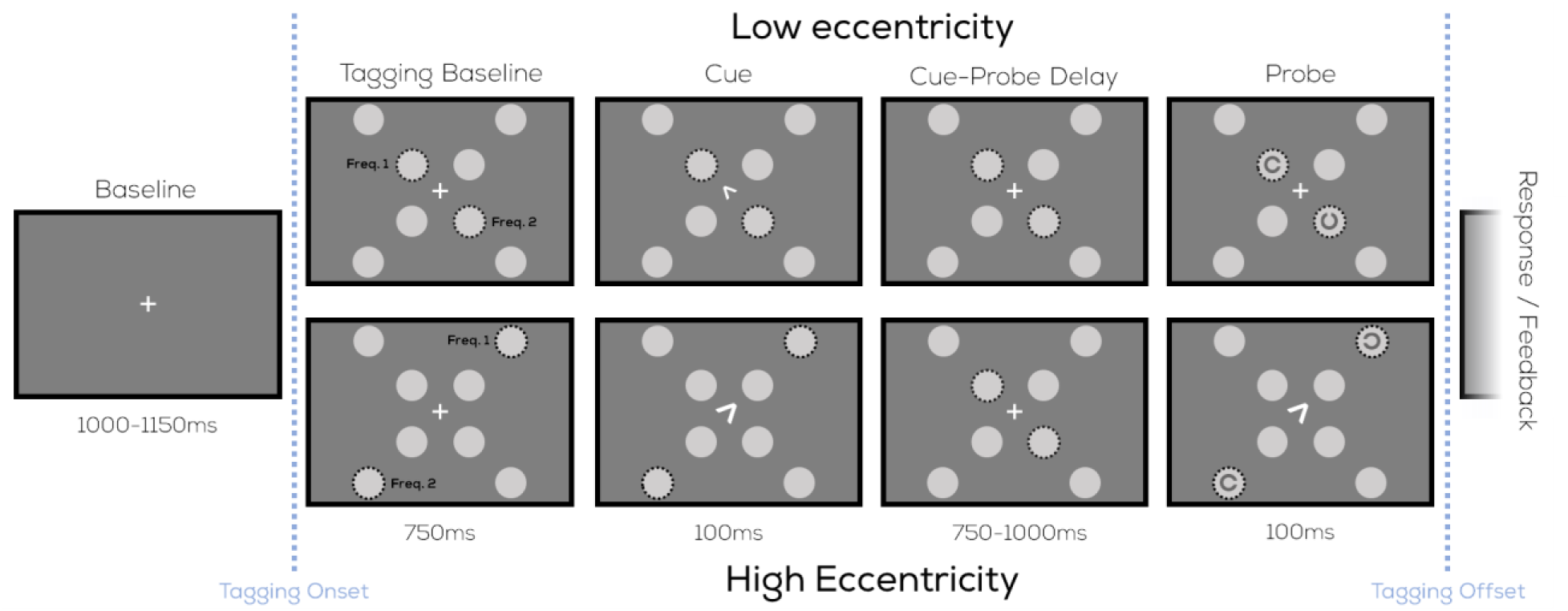
Task design. Participants reported the orientation of a target presented at one of eight locations as indicated by a spatial cue. Eight circles were presented around fixation, and a diagonally placed pair of circles was tagged at 60.0Hz and 65.5Hz (see *Tagging Manipulation*) throughout the trial. An arrow cue specified which of the eight locations the target will appear at. This cue was either **Top**: small, indicating that the near circle was cued, or **Bottom:** large, indicating that the far circle was cued. A target and distractor were presented at the tagged locations. Participants indicated the orientation of the target (facing left, right, up, or down).

At the start of the trial, the fixation cross was presented for a variable duration of 1-1.15s, followed by the arrangement of eight circles, of which a random (diametrically opposite) pair was flickered at 60.0Hz and 65.5Hz (0.75s). This was followed by an arrow cue (0.1s) that pointed to one of the two tagged circles, informing participants which location will be randomly cued on that trial. The size of the arrow cue was used to disambiguate between the low-eccentricity (smaller arrow cue) and high-eccentricity (larger arrow cue) circles in each diagonal direction. The cue was followed by a randomly uniform stimulus delay interval (0.75–1s), after which two Landolt-C stimuli were displayed (0.1s): one at the cued location (the target) and one at the diametrically opposite location at equal eccentricity (the distractor). Both locations were tagged. Participants were instructed to report the location of the aperture within the target Landolt-C (i.e., its orientation), which could be facing right, left, up, or down (responses were provided with the corresponding arrow key). The tagging was maintained until the end of the response period. Participants could respond up to 2s after the target onset. If no response was provided within this window, or if an incorrect button was pressed, the trial ended and participants were shown “incorrect” feedback (centrally presented; 0.25s). If the correct key was pressed, participants were shown “correct” feedback (0.25s). Each participant completed 900 trials (15 blocks of 60 trials each). The pair of circles that was tagged (4 possible pairs), the target location (2 possible locations within each pair), and the orientation of the target stimulus (4 possible orientations) were presented equally often (individually, not counterbalanced) and in a different random order for each participant.

### Stimuli

The screen background was kept at the same dark grey color throughout the experiment (i.e., 23% grayscale), visibly darker than the tagged stimulus regions. Inner circles were positioned at a diagonal eccentricity of 4.5 dva, and outer circles at 8.1 dva, both offset equally in the horizontal and vertical directions from fixation. All circles were identical in size (r = 1.6 dva) and visual appearance. The circles that were not tagged, were displayed with medium grey (50% grayscale; mean luminance of the flicker) to ensure that they appeared the same as the tagged stimuli regions. A white fixation cross composed of two perpendicular white lines (length = 0.65 dva, line width = 0.11 dva) was presented in the center of the screen on each trial. The fixation cross stayed on the screen except for when it was replaced by the arrow cue and the feedback image (Tick = ‘correct’; X mark = ‘incorrect’). The arrow cue was constructed from two white line segments (line width = 0.11 dva) extending from the fixation point in one of the four diagonal directions. For arrows pointing to the near circles, each segment had a length of 0.32 dva and for arrows pointing to the far circles, each had a length of 0.65 dva. The target and the distractor were both black Landolt Cs (C-shaped rings). The inner Landolt-C was 1.14 × 1.14 dva, and the outer Landolt-C was approximately 1.32 × 1.32 dva (width × height). Both Landolt-Cs were rotated to face one of four cardinal directions (0°, 90°, 180°, or 270°). Stimulus opacity was adaptively adjusted using a Bayesian staircase (Farell & Pelli, 1999) to maintain the accuracy on the task at 70% (PsychtoolBox QUEST algorithm, β = 3.5, Δ= 0.01, γ = 0.25). Separate staircases were run for the inner and outer target locations.

### Tagging Manipulation

In this study, Rapid Invisible Frequency Tagging (RIFT) was used to evoke a unique, frequency specific response that reflects the degree of visual processing of specific locations on the screen. To this end, a pair of diagonally opposite circles (randomly selected on each trial) was tagged until a response was recorded. Two tagging frequencies were used here; 60.00Hz (i.e., 360/6), and 65.4545…Hz (i.e., 360/5.5). Throughout the manuscript, we round these frequencies to 60.0Hz and 65.5Hz. The tagging process involved sinusoidally modulating the screen luminance at two circle locations with these fixed frequencies. Given the 360Hz refresh rate, for the 60.0Hz tag this required that we sample 6 equally spaced points on one cycle of a sine wave and assign these values as luminance levels to 6 consecutive frames of the tagged stimulus display. This set of 6 luminance values was then repeated over consecutive frames which formed 60 cycles of luminance oscillations in a 1 second period. Similarly, the 65.5Hz tagging envelope was computed by sampling 11 equally spaced points on two cycles of a sine wave (producing, on average, 360Hz/65.5Hz = 5.5 points per cycle). We did not implement any transparency masks to produce smooth edges of the tagging regions, or any ramping up/down of the tagging amplitudes over time.

The two frequencies (60.0Hz and 65.5Hz) were pseudo-randomly assigned to either the cued or the uncued circle of the diagonal pair on each trial. The sinusoids were always at the same phase at the moment of arrow cue onset. Temporal precision of the displayed stimuli was continually recorded using PsychToolBox’s Screen(‘Flip’) command. To account for any missed frames, an online correction was implemented that adjusted the phase of the tagging waveform forward to maintain its original phase trajectory when this command detected missed frames (mean percentage of trials per participant with one or more missed frames = 17.72%, SD = 19.84%, median = 11.88%). Note that it is a limitation of the current hardware setup that we were unable to achieve consistent frame display at 360Hz, with such a large proportion of missed frames. Rather than implement such an online phase correction, we would recommend for future work that researchers utilize the detailed hardware and software guidelines put forth in Dimigen et al., 2025.

### Protocol

Participants took part in a 2-hour session conducted at the Division of Experimental Psychology, Utrecht University. Before the experiment sessions began, participants received procedural information and gave informed consent. On the informed consent sheet, they were asked to report their date of birth, biological sex, and dominant hand. Once the EEG setup was complete, participants were seated at a fixed distance of 56 cm from the monitor, with their chins positioned on a chinrest. The experiment was verbally explained using a visual guide. The participants were instructed to maintain a fixed gaze on the fixation cross in the center of the screen. Participants were also informed that they might occasionally notice visual artifacts or flickering on the screen and were later asked whether they did or not. They were also informed that the transparency of the targets might vary across trials. After receiving these instructions and completing a practice block of 15 trials, participants proceeded with the main experiment, which lasted approximately 50 to 60 minutes. Compensation was awarded either in cash or in participation credits (see *Participants*), and the session was ended.

### Display Apparatus

Stimuli were presented on an LED-backlit LCD Alienware AW2524HF monitor (projected screen size: 54.5 × 30.5 cm (width × height)) (Dell Technologies, 2023), with a resolution of 1920 × 1080 pixels and a refresh rate of 360Hz. Beyond modifying the refresh rate to this particular setting, no changes to defaults (eg. brightness, contrast) were made on the monitor. Experimental code was written using PsychToolBox (Kleiner et al., 2007) in MATLAB (MATLAB, 2022).

### EEG Recording and Pre-processing

EEG data was recorded using a 64-channel ActiveTwo BioSemi system (BioSemi B.V., Amsterdam, The Netherlands) at a sampling rate of 2048Hz, with a Common Mode Sense (CMS) and Driven Right Leg (DRL) based online reference. To monitor vertical and horizontal eye movements, two extra electrodes were positioned above the left eye and on the outer canthus. Just before the experiment began, signal quality across all channels was checked using BioSemi’s ActiView software to ensure adequate recording quality. All data analysis was conducted using MATLAB’s Fieldtrip Toolbox (Oostenveld et al., 2011). The EEG data was first re-referenced to the average of all EEG channels (excluding poor channels, as determined by visual inspection, median = 13 channels per participant). The data was high-pass filtered (0.01Hz), then line noise and harmonics were removed using a DFT filter (50, 100, 150Hz). The data was segmented into trials ranging from 0.7 seconds before to 2.15 seconds after tagging onset. An ICA was performed on the segmented data to remove oculomotor artifacts (1 component removed per participant as per visual inspection), and trials with other motor artifacts or high noise (as per visual inspection) were removed (mean = 16.26%, SD = 12.20%, median = 14.61% of trials).

### RIFT Response: Coherence

Magnitude-squared coherence was used to determine the strength of the EEG response to RIFT frequencies of interest. Coherence is a dimensionless quantity ranging from 0 to 1 that quantifies the consistency of two signals in their frequency and phase alignment. High coherence values are achieved when two signals oscillate at the same frequency with a consistent phase difference across trials. Using coherence provides significant advantages in filtering out noise, since only signals that both oscillate at the tagged frequencies (60.0Hz/65.5Hz) and maintain a consistent phase across trials influence the output. Coherence was computed between a reference wave (pure sinusoids at the corresponding 60.0Hz/65.5Hz frequency, sampled at 2048Hz) for a set of trials per channel and participant. The 2.85s trials were first bandpass filtered (±1/2.5Hz) at the frequencies of interest (60.0Hz & 65.5Hz) using a two-pass Butterworth filter (4th order, Hamming taper). The filtered time-series data was Hilbert transformed. This yielded the instantaneous magnitude, M_x_(t), and phase, Φ_x_(t), of the bandpass-filtered EEG signal. To compute time-varying coherence, the following equation was used:

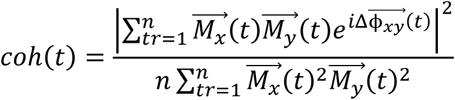

where M_x_(t) and M_y_(t) are the instantaneous magnitudes of the EEG signal and the reference sinusoid, respectively, and Φ_xy_(t) is the difference between their instantaneous phases across all n trials. We computed coherence for each channel individually. Then, for all plots except topographical, we selected the top six channels with the strongest coherence at 60.0Hz and 65.5Hz across all trials for each participant. All subsequent comparisons across experimental conditions were based on coherence traces averaged across these top six channels. Previous work (Arora, Gayet, et al., 2025; Arora et al., 2026) has shown that the number of selected channels has negligeable impact on relative coherence differences between experimental conditions. Coherence spectrograms were created to inspect the frequency specificity of the tagging signal by calculating coherence across frequencies ranging from 57Hz to 67.5Hz in 0.5Hz intervals. All subsequent comparisons across experimental conditions were based on traces averaged across the top six channels.

When making comparisons between cued and uncued RIFT responses, any attentional comparisons we make are never between the two diametrically opposite tagged locations on the same trial. Such comparisons would be vulnerable to hemispheric asymmetries in overall RIFT strength, and to differences between tagging frequencies, which could influence the resulting differences. Instead, we compare trials where a given tag was presented at a given location, and cued, to trials where the same tag was presented at the same location, but uncued. That is, our cued vs. uncued comparisons are always made within a location. While analyzing these attentional comparisons, we also chose to include all trials, both correct and incorrect. This is because here, we staircased the transparency (=visibility) of the Landolt-C targets in order to produce a ∼70% accuracy level. This means that responses would often have been incorrect simply due to the visibility of the Landolt-C being so low that even in the event of a successful shift of attention, the target was barely below the threshold of discriminability. In this case, an incorrect report would not (necessarily) reflect a failure to shift attention, but rather a failure to determine the target orientation. This is why we also include incorrect trials. However, we also replicate the same effects as those of our main attentional comparisons when using only correct trials in Supplementary Figure 2.

### General Statistical Analysis

A cluster-based permutation test was used to compare condition-wise attentional differences in coherence over time (Maris & Oostenveld, 2007). First, coherence traces were averaged over the six channels with the strongest RIFT coherence at 60.0/65.5Hz for each participant, yielding a single coherence trace over time for each condition at each frequency. To investigate the differences across attentional conditions, the uncued condition trace was subtracted from the cued condition trace for each frequency and participant. We then applied a cluster permutation test on the entire trial duration (0.7s before to 2.15s after tagging onset) to assess statistical significance (Maris & Oostenveld, 2007). A one-sample t-test was performed at each individual time point on this difference trace to identify intervals that were significantly different from zero (p < 0.05). Clusters were defined as sequences of consecutive significant time points, and only those spanning at least 10ms were retained. For each cluster, a cluster-level test statistic was computed by summing the corresponding t-values (t-mass). To construct a reference distribution, we randomly flipped the sign of each participant’s difference trace to preserve temporal autocorrelations while removing consistent condition effects. This procedure was repeated 5000 times, and the maximum t-mass from each permutation was stored to form the reference distribution. Finally, we assessed whether the t-masses of any observed clusters exceeded 95% of the null distribution, and clusters exceeding this threshold were considered statistically significant. Coherence values were initially compared between cued versus uncued locations separately for each tagging frequency, and results were later averaged across frequencies. To visualize the variability of coherence traces across participants, we used 95% confidence intervals after applying a correction to account for cross-participant variation in the overall RIFT signal (Cousineau, 2005).

The above described cluster permutation test was also used to identify significant tagging responses when computing coherence based on different tagging locations. In this case, instead of using a difference trace computed between attentional conditions for each participant, a difference trace was computed between the post- and pre-tagging onset periods across frequencies. Lastly, when comparing time-averaged coherence values from different spatial conditions (low-vs. high-eccentricity, lower vs. upper hemifield, left vs. right hemifield), we used two-sided paired-sample Bayesian t-tests with default priors using the bayesFactor package in MATLAB (Krekelberg, 2025).

## Results

Behavioral results from the experiment showed that participants performed the task successfully. The overall performance was above chance (mean = 72.29%, 95% CI: [71.61%, 73.50%]), matching the 70% accuracy set by the staircase procedure (see Stimuli). Accuracy on high-eccentricity trials (mean = 73.48%, 95% CI: [72.28%, 75.21%]) and low-eccentricity trials (mean = 71.13%, 95% CI: [70.56%, 72.61%]) was significantly different (p = 0.0024; bootstrapped test of mean differences), however, both were similar in magnitude. All further analyses shown here were conducted using both correct and incorrect trials (see “RIFT Response: Coherence” for the reasoning behind this choice and Supplementary Figure 2 for control analyses that replicate our main findings with only correct trials), however, we also repeated our analyses on the correct trials alone and observed comparable statistically significant attentional effects following cue presentation.

### Tagging manipulation on Alienware AW2524HF monitor (refresh rate = 360Hz) evokes a RIFT response at the frequencies of interest

First, we verified that our tagging manipulation successfully evoked an oscillatory response in the EEG signal. Coherence showed a clear peak at the tagged frequencies 60.0Hz and 65.5Hz (Figure 2A), demonstrating that we successfully captured two simultaneous RIFT tags. In line with previous RIFT studies, coherence at the lower frequency (60.0Hz; mean = 0.014, 95% CI: [0.009, 0.028]) was significantly larger in magnitude (p = 0.009; bootstrapped test of mean differences) than coherence at the higher frequency (65.5Hz; mean = 0.007, 95% CI: [0.004, 0.017]). Coherence was highest around central/parietal channels (Figure 2B), unlike previous RIFT studies (Arora, Gayet, et al., 2025; Bouwkamp et al., 2025; Dimigen et al., 2025; Spaak et al., 2024; Zhigalov et al., 2019), where coherence was highest around parietal/occipital channels.

**Figure 2:**
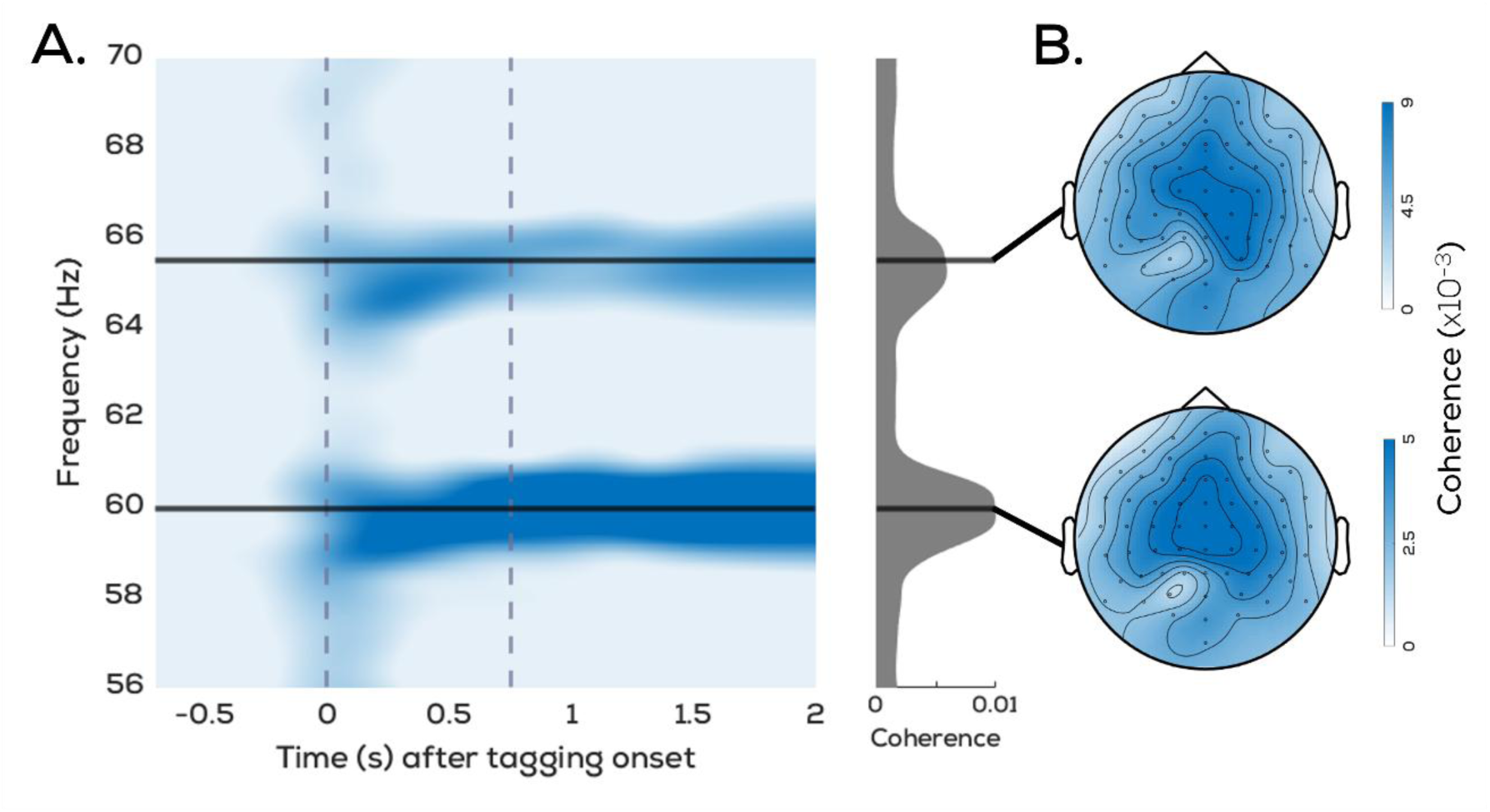
Average RIFT response. **A**. Coherence averaged across participants and top channels (see *RIFT Response: Coherence*) showing clear peaks at 60.0Hz and 65.5Hz following flicker onset. Vertical dashed lines indicate the tagging (t=0s) and cue (t=0.75s) onset. Time-averaged distributions in grey are based on the full spectrogram time window (t=-0.7s to t=2s). **B.** Topographical distribution map of average RIFT coherence over the interval during which tagging was on (t=0s to t=2s).

### RIFT responses vary across different locations in the visual field

**Figure 3:**
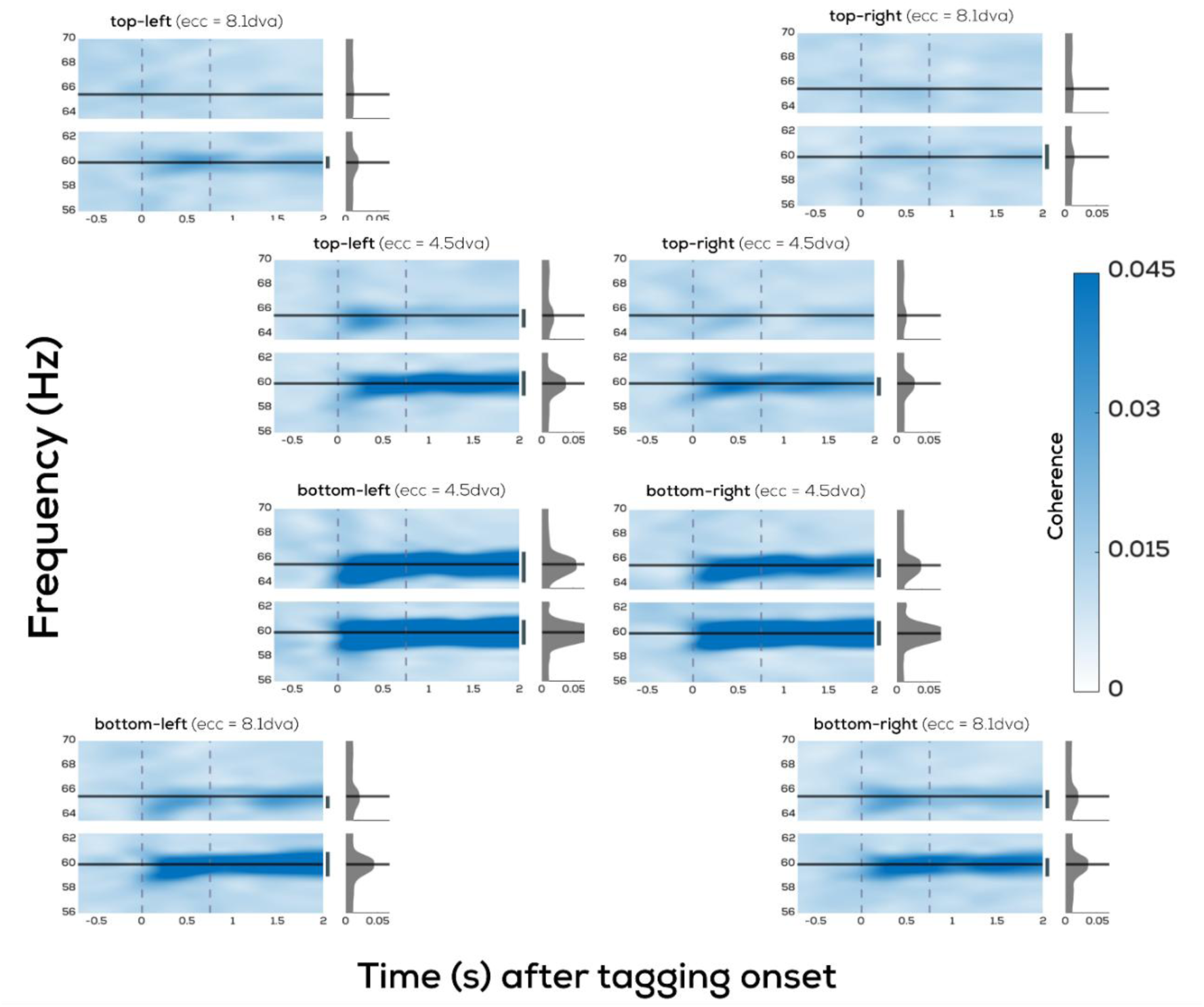
Average RIFT response by tagging location. Spectrograms of mean coherence split by tagging location. Panels are positioned according to the arrangement of tagged circles in the experimental display (see Figure 1). The black horizontal lines are drawn at 60.0Hz and 65.5Hz. Vertical dashed lines indicate the tagging (t=0s) and cue (t=0.75s) onset. Significance bars reflect the results of cluster based permutation tests (see *General Statistical Analysis*).

We investigated the extent to which RIFT responses differ for different locations in the visual field (see Figure 1 for task design) to determine whether more peripheral tagging regions can also evoke a clear RIFT response. We picked up on significant 60.0Hz tagging responses from all eight locations used in this study (Figure 3). Significant 65.5Hz responses were observed from all locations in the lower hemifield, but only the low-eccentricity-left tag in the upper hemifield (Figure 3). In line with existing studies with multiple tagging locations (Minarik et al., 2023; Wang et al., 2026), we found significantly higher tagging responses overall from low-eccentricity tags compared to high-eccentricity tags (BF10: 95.62; Figure 4A), and from the lower hemifield compared to the upper hemifield (BF10: 160.85; Figure 4B). The fact that upper hemifield responses are consistently reduced compared to lower hemifield responses (Minarik et al., 2023; Wang et al., 2026), and the fact that higher frequencies consistently evoke lower responses (Arora, Gayet, et al., 2025; Gulbinaite et al., 2019; Minarik et al., 2023; Wang et al., 2026), makes it unsurprising that the cases in which the lowest tagging responses were observed here were those of the upper hemifield tags at 65.5Hz. We did not observe a difference between the tagging responses from left and right tagging locations (BF10: 0.88; Figure 4C). Lastly, the topographical spread of coherence across the different tagging locations (Figure 5) suggests that the RIFT response in more central channels comes mainly from the tagging locations in the lower hemifield.

**Figure 4:**
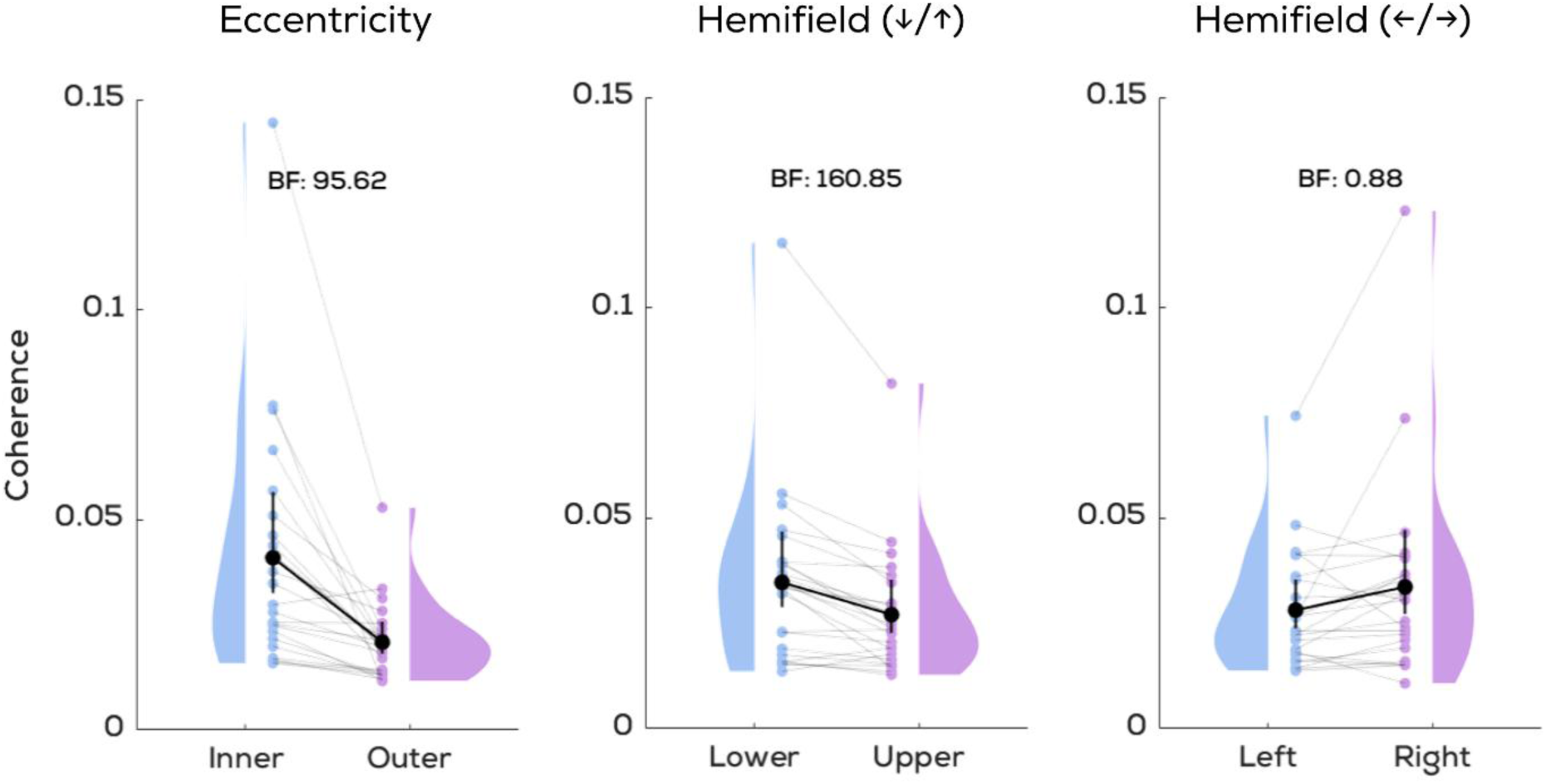
Pairwise comparisons of RIFT responses by location. Mean coherence compared between various spatial conditions (low vs. high eccentricity, lower vs. upper hemifield, left vs. right hemifield) using bayesian t-tests. RIFT responses are significantly higher from low as compared to high eccentricity tags, and from lower as compared to upper hemifield tags. Shaded regions reflect kernel density estimates, and black bars reflect 95% CIs.

**Figure 5:**
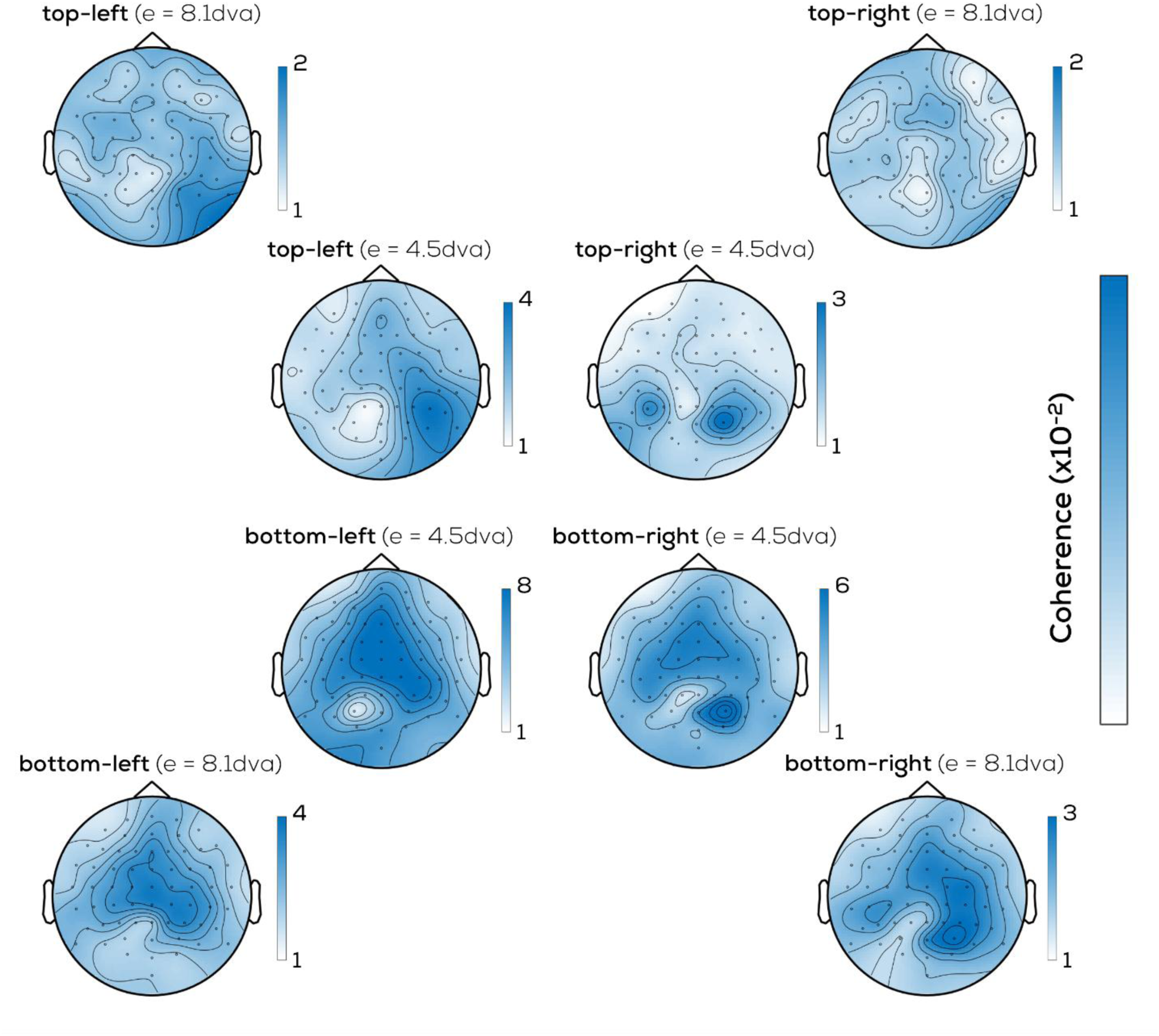
Average RIFT topography by tagging location. Topographical distribution of average RIFT coherence during the baseline tagging interval, averaged over 60.0Hz and 65.5Hz. Panels are positioned according to the arrangement of tagged circles in the experimental display (see Figure 1). Note that individual topoplots follow separate colour axis limits.

### Attention modulates the RIFT Response

We compared coherence traces between cued and uncued locations to investigate attentional modulations of the RIFT response (Figure 6A). A cluster permutation test indicated a significant difference between cued and uncued items soon after cue onset (p = 0.0002). This demonstrates that a RIFT-monitor setup can measure attentional modulations of the RIFT response before (during the spatial attentional shift) and after the appearance of the target stimulus. Lastly, we confirmed that this attentional modulation was not driven only by the tagging regions in the lower hemifield which had evoked an overall higher RIFT response (Supplementary Figure 3).

**Figure 6:**
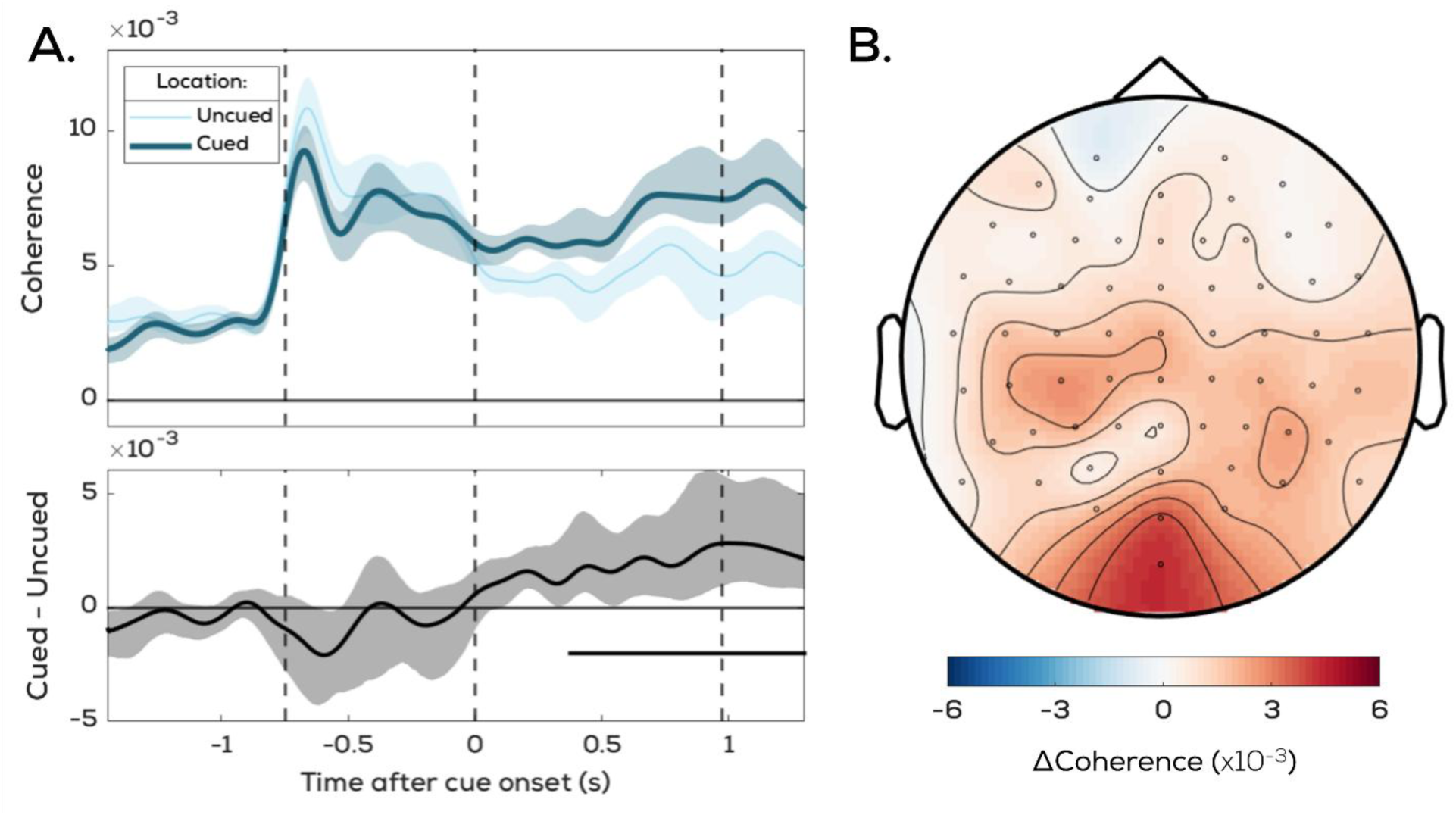
**A. Average RIFT coherence from cued versus uncued locations.** Coherence traces reflect signals from locations corresponding to cued and uncued tags, with their difference below. Solid black line marks significance as per the cluster permutation test. The vertical lines represent the tagging, cue, and target onsets, respectively. The target onset is marked at the midpoint between the minimum and maximum delays (0.85s –1.1s) following the arrow cue offset. **B. Topographical distribution of the attentional modulation.** Average difference between coherence from cued and uncued tags (averaged over the interval significant at the group level).

The topographical distribution of the difference in RIFT coherence between cued and uncued item locations, collapsed across both flicker frequencies and averaged over the post-cue interval, shows a positive effect predominantly over parietal-occipital channels (Figure 6B). Though this spatial distribution of attentional modulations is a marked shift from the overall coherence topographies (Figure 2B) as it shows a more posterior distribution, it aligns with previous RIFT work (Arora, Gayet, et al., 2025; Dietz et al., 2025).

## Discussion

This study implemented Rapid Invisible Frequency Tagging (RIFT) along with an attentional cueing task on a 360Hz refresh rate consumer-grade monitor. We show that attentional modulations of the RIFT response can be captured on a monitor-based display setup. Our findings present an essential extension to recent work by Dimigen et al. (2025), who showed that a 480Hz refresh rate monitor can be used to evoke RIFT responses at 60.0Hz and 64.0Hz, however, without testing if these responses are modulated by attention. This is a notable advancement in the field of RIFT, as all existing RIFT research prior to this has utilized the PROPixx projector, which is a relatively expensive piece of specialized display hardware (∼30x the cost of the monitor used here). We show here that a similar monitor setup (Alienware Gaming Monitor AW2524HF at 360Hz refresh) can produce frequency-specific neural responses at 60.0Hz and 65.5Hz and, more importantly, that it can measure modulations to these tagging responses by covert attention via a simple cueing paradigm.

We show that upon cueing participants to a location on the screen for the report of an upcoming target, the tagging response is significantly higher for cued as compared to uncued locations. This attentional modulation can already be observed prior to target onset. Thus, we show that even when using a consumer-grade display setup, the evoked RIFT response is sensitive enough to capture modulations to early visual processing induced by covert shifts of attention.

Although prior RIFT-monitor work measured weak tagging responses when tagging peripherally (Dimigen et al., 2025), we show here that peripheral tagging on a monitor, at least at 60.0Hz, is both reliably observable as well as able to reflect attentional modulations. However, it is worth noting that our tagging was less eccentric (4.5 dva and 8.1 dva) than the peripheral tagging in Dimigen et al. (2025) (12 dva), which is the likely cause of the difference in tagging strength. The eccentricity range used here is generally sufficient for a vast range of cognitive experiments. Beyond this, our observations regarding how tagging responses vary across the visual field mirror previous work that has investigated this more thoroughly (Minarik et al., 2023). Specifically, the findings we replicate here are that the lower visual field evokes a higher RIFT response, that more peripheral tags evoke a lower RIFT response than less peripheral ones, and that higher frequencies evoke lower RIFT responses.

A key consideration in RIFT experimentation is the invisibility of the tagging. Since this is generally assumed to be a function of refresh rate (i.e., higher refresh rates are less susceptible to visibility of the tagging), this is a key concern with monitor setups such as the 360Hz setup used here as compared to the 1440Hz refresh rate of the PROPixx projector. Although no participant perceived the flicker continuously, they all reported being aware of some screen changes -likely the visual glitches on the screen caused by missed frames. Though we did not empirically test for tagging visibility here, these reports are similar to those of Dimigen et al. (2025). Further research should be conducted to verify whether this monitor setup reliably achieves invisible frequency tagging before using it to tag locations that are perceptually indistinguishable from background space, where the flicker is more susceptible to being visible. However, when tagging visible stimuli (rather than empty background space), as was the case here, RIFT implementations paired with monitor setups appear to be promising low-cost alternatives to using expensive projectors. This is an important development for M/EEG researchers studying visual cognition with computerized tasks: now, stimuli in existing tasks can be tagged to obtain an extra time-varying measure of attentional allocation without needing an expensive device dedicated to RIFT studies. Whether the behaviour of attention itself is being studied, or attention is a possible confound in the task, having an additional measure at little cost to the experimental design can be an immense benefit to cognitive tasks.

Another consideration when using frequency-tagging approaches to measure the allocation of spatial attention is eye movements. Since it is well-established that even covert shifts of spatial attention are accompanied by small (<0.5dva) eye movements towards the attended location (Engbert & Kliegl, 2003; Hafed & Clark, 2002; Van Ede et al., 2019), it is possible that eye movements bring the cued tag closer to the fovea and in doing so drive the increased tagging response from this location. However, several previous studies with similar tasks have conducted analyses with various approaches showing that these effects do not correlate; i.e., modulations to RIFT responses can be observed independently of gaze position changes (Arora et al., 2026; Arora, Gayet, et al., 2025; Arora, Van der Stigchel, et al., 2025; Wang et al., 2026). Though gaze position data was not recorded in the current study, an additional analysis using electrooculography (EOG) data also shows this distinction here (Supplementary Figure 1). This EOG analysis, combined with the previous literature, supports the claim that the RIFT modulations observed here stem from neuronal excitability rather than gaze position.

The definition of “uncued” tags in our tasks may influence the type of attentional modulation that we infer here. We specifically select the stimulus regions diametrically opposite to the cued tags as “uncued” tags. What we compare with this design, after balancing for tagging frequency and location, are RIFT responses when a region was cued and when the region diametrically opposite to it was cued. We then infer attentional modulations by comparing the two and checking whether cued tagged regions display stronger RIFT responses than uncued tags, which they consistently do. This means, however, that we cannot distinguish here how this difference arises. That is, we cannot distinguish whether attention is implemented only by boosting visual processing at the cued location, or, by doing so and also suppressing visual processing at the diametrically opposite location of the visual field. To distinguish such mechanisms, future studies should ideally tag a greater number of regions on the screen. This could be by using, for example, phase in combination with RIFT (Arora et al., 2026; Spaak et al., 2024).

Here, we observed notably different topographies of the RIFT response as compared to previous studies, with a more prominent central cluster rather than the typical occipito-parietal clusters. However, we refrain from interpreting this as necessarily reflecting a different neuronal source. In general, it is very difficult to link EEG scalp projections to underlying sources, because of the complexities associated with the dipoles that give rise to these EEG projections. The channels which show the strongest responses need not be directly above the underlying source. We believe the biggest factor in determining the topographical pattern observed is stimulus characteristics (size and position). For example, although it is expected (and common) to see a contralateral RIFT response when lateralized tagging stimuli are presented, we have previously observed both contra- and ipsilateral peaks (Wang et al., 2026; see Supplementary Figure 7) from one tag. This means that even slight changes in size, eccentricity, and polar angles can produce unexpected topographies. We also used relatively small stimuli here; whereas larger stimuli may produce a more “average” topography by eliciting responses across a larger cortical region, smaller stimuli will produce more specific and targeted patterns of activity. A qualitative observation of Figure 2 (and the most strongly responsive regions -inner bottom-left and bottom-right -in Figure 5) shows a distinct absence of a RIFT response at the channels we would most expect it, surrounded by two clusters more centrally and more dorsally. This pattern is consistent with a dipole centered around the expected posterior channels, projecting two clusters centrally and dorsally. It is most likely that the specific eccentricity and angles we used here resulted in a pattern where a dipole is centered around the same area as previous RIFT responses, but is simply oriented differently, projecting two clusters more centrally and more dorsally.

These findings offer sufficient evidence in support of a 360Hz refresh-rate consumer-grade monitor as a display apparatus compatible with RIFT, forming a far more affordable alternative to the projector setups traditionally used in RIFT research. Our results show that RIFT with a monitor implementation can successfully capture neural responses to multiple rapidly flickering stimuli, as well as modulations to these responses by covert spatial attention. By doing so, we demonstrate that consumer grade monitors are a valid choice for implementing invisible frequency tagging. We hope that this study will serve as a basis for researchers studying attention or using tasks where visual attention may be a possible confound, who can now use relatively inexpensive monitor setups to gain an imperceptible tracker of attention with only minor adjustments to their tasks.

## Declarations

### Funding

This research was funded by a VICI Grant from the Netherlands Organization for Scientific Research to Stefan Van der Stigchel.

### Competing Interests

The authors declare no competing interests.

### Ethics Approval

The study was conducted in accordance with the protocol approved by the Ethics Committee of the Faculty of Social and Behavioral Sciences at Utrecht University with reference number 863732 –HOMEOSTASIS (approved on 18/03/2020).

### Consent to participate

Informed consent was obtained from all individual participants included in the study.

### Consent for publication

Informed consent was obtained from all individual participants included in the study. The authors affirm that human research participants provided informed consent for publication. No identifying information is included.

### Availability of Data

Preprocessed and raw EEG Data is available at https://osf.io/d8vp5/overview

### Code Availability

Experimental code and analysis code is available at https://osf.io/d8vp5/overview

### Author Contributions

Conceptualization, K.A., G.C., L.K., S.V.d.S., S.G., S.C.; data curation, K.A., G.C., L.D.; formal analysis, K.A., G.C.; funding acquisition, S.V.d.S.; investigation, K.A., G.C.; methodology, K.A., G.C.; software, K.A., S.C.; supervision, L.K., S.V.d.S., S.G., S.C.; visualization, K.A., G.C.; writing—original draft preparation, K.A., G.C.; writing—review and editing, K.A., G.C., L.D., L.K., S.V.d.S., S.G., S.C.

## Supplementary Information

**Figure S1:**
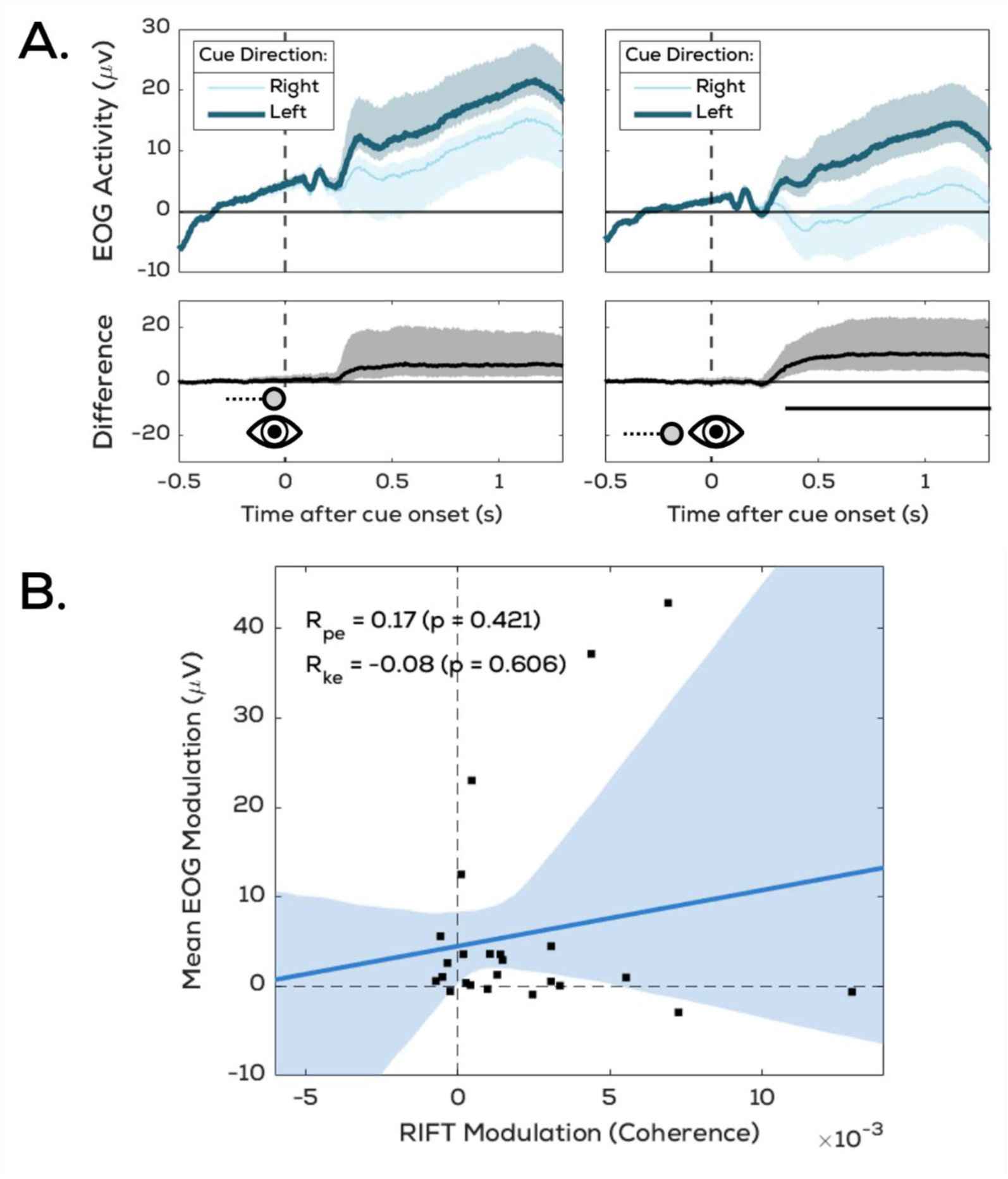
EOG-derived gaze activity does not correlate with attentional RIFT modulations. **A.** Shifts of spatial attention, even when covert, are usually accompanied with small (<0.5dva) biases in gaze position through microsaccadic eye movements. Although no eyetracking data was recorded here, we can nonetheless observe these cue-induced gaze biases here through the electrooculography (EOG) response. This is shown here with significant differences in the response from the EOGs above the left eye (Left) and on the outer canthus (Right) when comparing trials cueing the left and right directions. **B.** However, this cue-induced modulation in EOG voltage does not correlate with the cue-induced modulation in the RIFT response. We averaged the participant-wise differences from both EOG electrodes in the post-cue period (from 0s to 1.5s after cue onset) to obtain one EOG modulation value per participant. We similarly averaged the RIFT cued-uncued difference traces (see Figure 6) per participant to obtain one RIFT modulation value per participant. We performed both parametric (Pearson coefficient) and non-parametric (Kendall Tau coefficient) correlations between these which showed that they are independent of each other.

**Figure S2:**
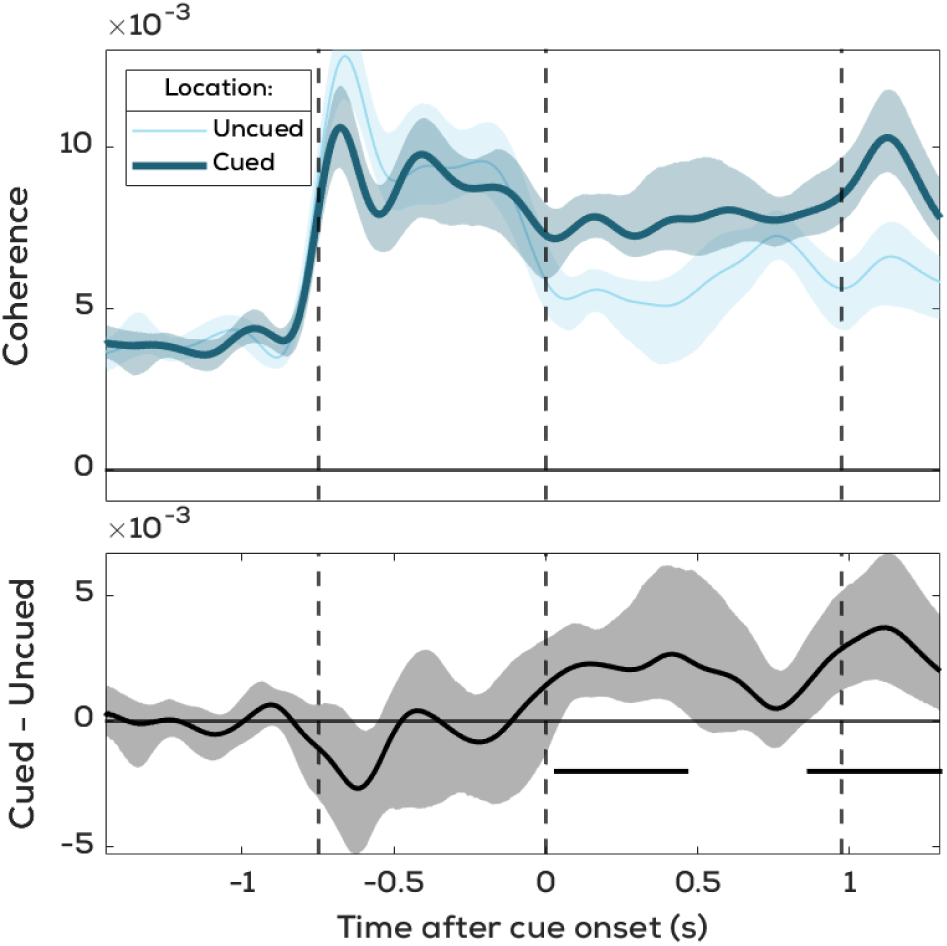
Attentional modulations are also observed when only using correct trials. Figure details are the same as Figure 6 in the manuscript.

**Figure S3:**
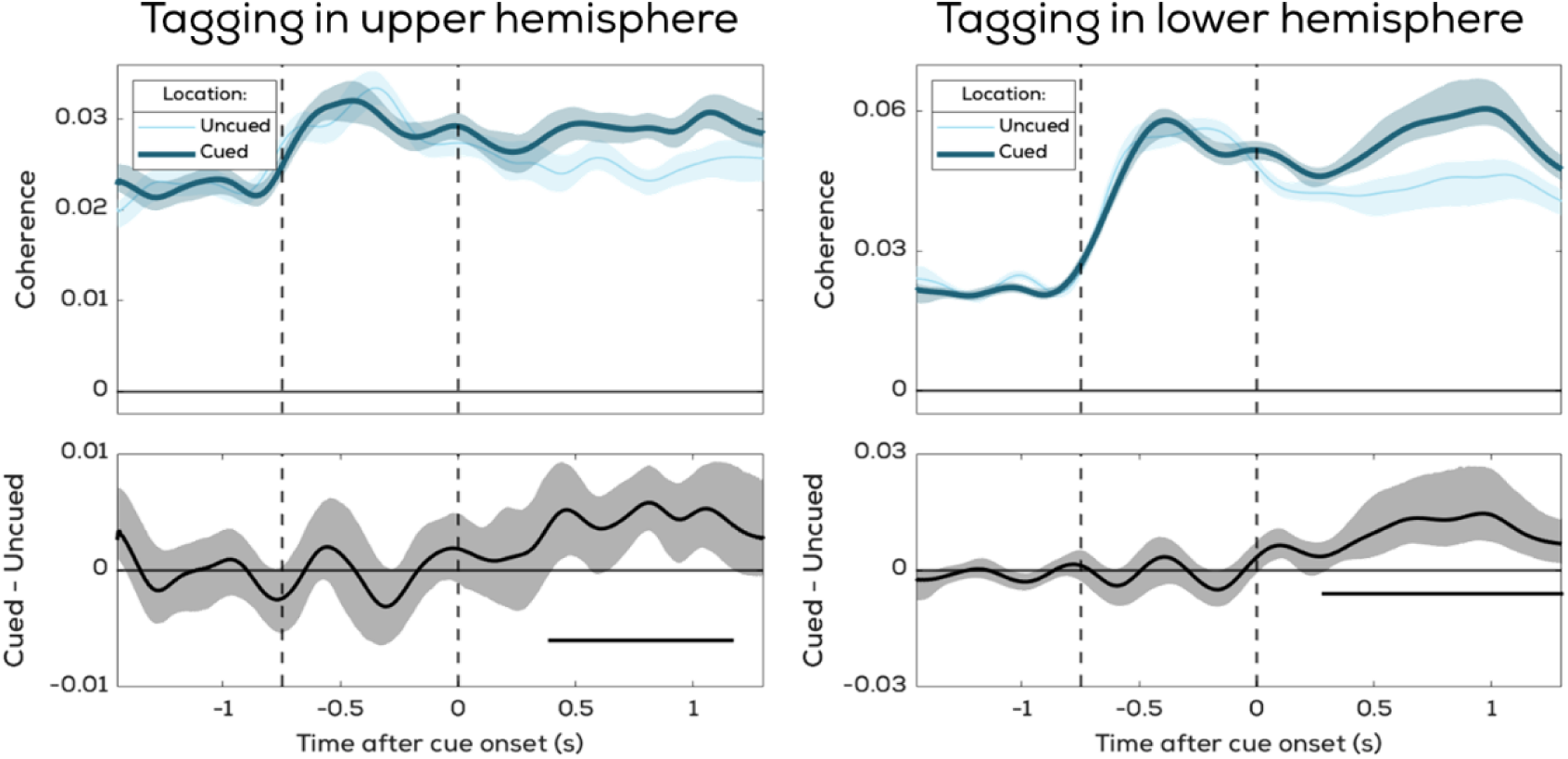
Attentional effects are present regardless of whether tags are presented in upper or lower hemifield. In the manuscript, we show that the upper hemisphere has a markedly weaker overall RIFT response. However, in Figure 6 we present attentional effects based on data from all tagging locations. Here, we show that the attentional modulation in Figure 6 is not driven only by lower hemisphere tags. Rather, both hemispheres show significant tagging modulations.

## Notes

### Competing Interest Statement

The authors have declared no competing interest.

### Summary of Updates

Correction to spectrogram showing RIFT responses per tagging location; Updated Figures; New Supplementary Material

